# Dynamics of cortical degeneration over a decade in Huntington’s Disease

**DOI:** 10.1101/537977

**Authors:** E. B. Johnson, G. Ziegler, W. Penny, G. Rees, S. J. Tabrizi, R. I. Scahill, S. Gregory, the TRACK-HD and TrackOn-HD investigators

**Author notes:** These authors contributed equally to this work (E. B. Johnson, G. Ziegler). Correspondence Eileanoir B. Johnson Huntington’s Disease Centre UCL Queen Square Institute of Neurology University College London United Kingdom Gabriel Ziegler Institute of Cognitive Neurology and Dementia Research Otto-von-Guericke-University Magdeburg Leipziger Str. 44 39120 Magdeburg Germany Phone: +49 / 391 67 250 54.

## Abstract

The neurodegenerative process is typically slowly progressive and complex. While simple models of neurodegeneration suggest that brain changes progress at a near constant rate, previous research shows regional variation within the temporal progression of atrophy, indicating that over the course of neurodegeneration, different regions may undergo changing rates of atrophy. Characterization of long-term dynamic brain changes in neurodegeneration requires both extensive longitudinal MRI datasets and an advanced modeling framework. Until recently, both of these elements were not available. Here, we implement a novel dynamic systems approach to infer patterns of regional progression spatially and temporally in a unique longitudinal dataset with up to seven annual individual brain scans per participant from 49 Huntington’s Disease (HD) gene-carriers. We map participant-and group-level trajectories of cortical atrophy in HD using a decade of data that encompasses motor symptom onset and, for the first time, show that neurodegenerative brain changes exhibit complex temporal dynamics of atrophy with substantial regional variation in progressive cortical atrophy. Some fronto-occipital cortical areas show an almost constant rate of atrophy, while medial-inferior temporal areas undergo only minor change. Interestingly, cortical sensory-motor areas were found to show a noticeable acceleration of atrophy following HD diagnosis. Furthermore, we establish links between individual atrophy and genetic markers of HD (CAG repeat length), as well as showing that cortical motor network changes predict subsequent decline in task-based motor performance, demonstrating face-validity of the model. Our findings highlight the complex pattern of dynamic cortical change occurring in HD that can help to resolve the biological underpinnings of HD progression.

## Introduction

Characterizing the trajectory of cortical atrophy is necessary for a mechanistic understanding of neurodegeneration. Simple models suggest that neurodegenerative changes follow a constant rate, however, brain areas may atrophy differently over the course of neurodegeneration^1–3^, and it is possible that rates of atrophy change over time. Discovering such possible regional accelerations and decelerations of cortical atrophy, that could reveal insights into the nature and mechanistic underpinnings of neurodegenerative disease, requires comprehensive spatio-temporal mapping of long-term brain changes. Until now, characterizing such dynamic patterns has been limited by the availability of modeling framework coupled to cohort data with multiple timepoints that could quantify the temporal evolution of patterns of long term brain atrophy.

Here, we solve this issue using a longitudinal data that spans a decade of disease progression to quantify the varying process of cortical atrophy in a well-characterized neurodegeneration cohort using a novel dynamic modeling technique. Our model offers a powerful new approach for defining brain changes by constructing individual participant-level dynamic models of atrophy over time, which are then used to identify group-wise trajectories of disease progression both spatially and temporally^4^. By specifying the temporal progression of cortical atrophy within all brain regions simultaneously^5^, we can characterize both total atrophy over a time period, plus changes (accelerations and decelerations) in rates of atrophy. These trajectories may be influenced by volumetric change in neighboring regions and/or external factors, such as genetic influences, which may speed up neural atrophy over time.

We used this model to characterize the changing trajectories of cortical atrophy in Huntington’s Disease (HD) over a 10 year period around the point of clinical disease onset^5^. HD is a progressive neurodegenerative disease, characterized by a triad of cognitive, psychological and motor symptoms^6^. Given the certainty of disease onset in those who inherit the HD gene, brain changes can be tracked from the early asymptomatic stage through to clinical motor disease onset. Atrophy in HD occurs in striatal regions many years prior to symptom onset with white matter (WM) and cortical changes evident with increasing disease stage^7–11^. Trajectories for this cortical change have been hypothesised^12^, but previous longitudinal studies have only described change occurring between two timepoints, typically over one or two year periods^9,10,13,14^, and they do not offer a reliable or comprehensive understanding of how the rate or distribution of atrophy might unfold with disease progression^15^.

We used this novel technique^4,5,16^ to infer disease progression in a cohort of HD gene-carriers from the well-characterized, multisite, longitudinal TRACK-HD and TrackOn-HD studies^7,9,10,17,18^. Participants underwent diagnostic conversion from pre-HD to manifest HD within a 10 year period surrounding motor diagnosis, a critical time encompassing physiological and macrostructural brain changes. We analyzed up to seven individual annual MRI scans and a measure of motor performance. For each individual, we focused on volume measures from widespread cortical (and subcortical) brain regions at each available timepoint using longitudinal registration and customized segmentation tools optimized for this cohort^19^. We first measured the total atrophy within each region, followed by regional rates of atrophy during this time; initially characterizing constant rates of atrophy, but also identifying accelerations and decelerations of atrophy that can occur differentially across the cortex. We investigated the associations between this atrophy and CAG repeat length, and whether between-region patterns of atrophy could be detected. Finally, we linked cortical atrophy to behavioral symptoms, testing whether atrophy could predict longitudinal performance on a motor task.

## Results

### Sample

In order to assess HD progression, we exploit a longitudinal design including repeated scanning of 49 patients with 3-7 (mean: 5.84 SD: 1.63) individual annual scans over (last scan) follow-up time of 2-6 (mean: 5.94 SD: 1.62) years. Thirty of the 49 participants had a total of 7 annual scans. Patient demographics at timepoint of motor diagnosis in the analyzed cohort are shown in Table 1 (see methods for details on inclusion criteria). Details on the exact timing of all longitudinal MRI acquisitions are presented in Supplementary Figure 1.

**Table 1.**
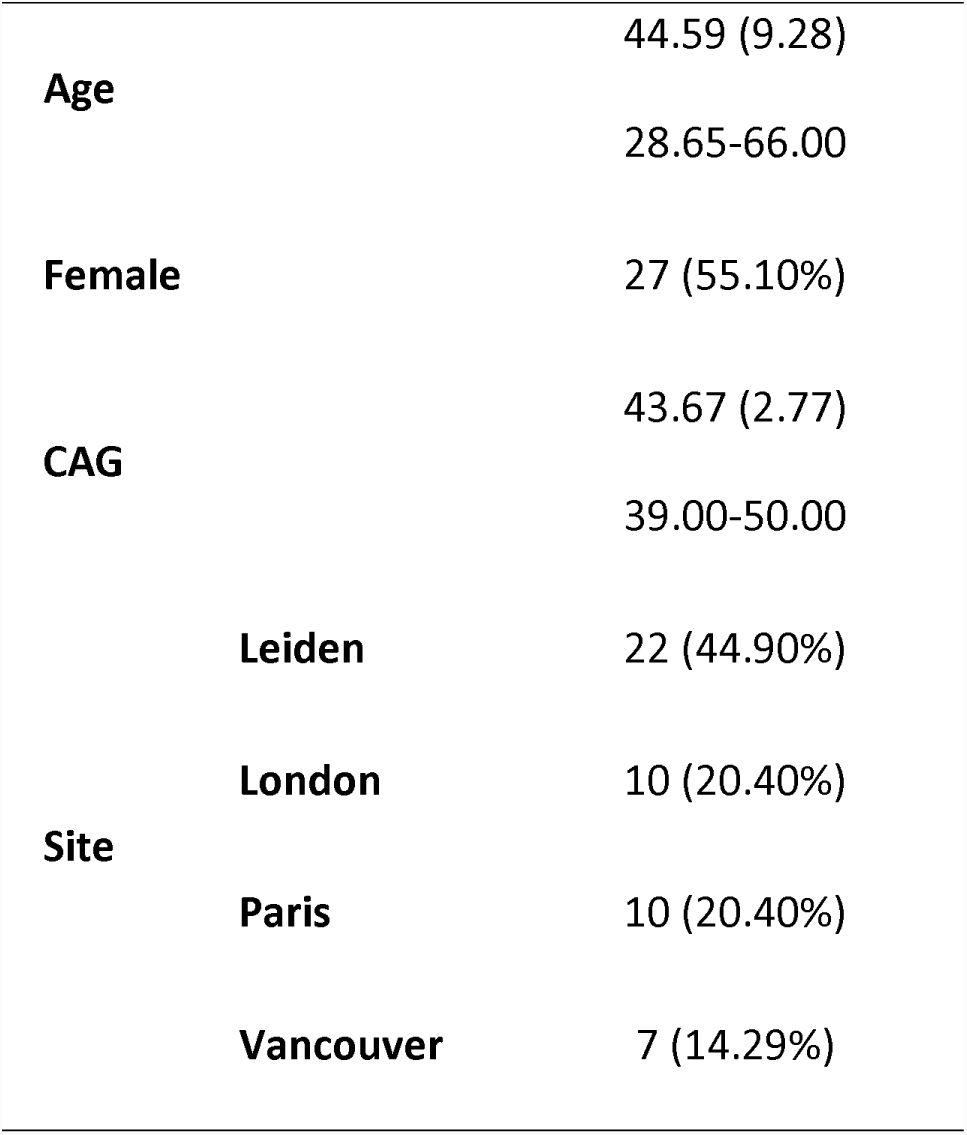
Demographics for the 49 participants included in this longitudinal multi-center study. The table shows mean (SD) and ranges, or N (%). As expected, HD carriers with longer CAG repeat length were found to have an earlier age when being clinically diagnosed (r=-0.85, p<e-13).

### Total atrophy over a decade is variable across the cortex

Examining changes over a decade of HD progression, the highest total volume reduction was observed in striatal regions; the putamen and caudate showing 21.11% and 19.68% loss respectively (Figure 1). We found volume changes in widespread cortical areas, especially in prefrontal, motor-and somatosensory regions, greatest total volume loss in supplementary motor cortex (11.66%), followed by frontal and precentral gyrus, superior parietal lobule, postcentral gyrus and inferior frontal gyrus, all showing 8.52-10.46% shrinkage. Global WM volume underwent an 8.25% reduction. These findings highlight that frontal and subcortical atrophy are both marked features of the long-term transition from preclinical to clinical HD, but suggesting a distinct pattern of spatio-temporal impairment.

**Figure 1.**
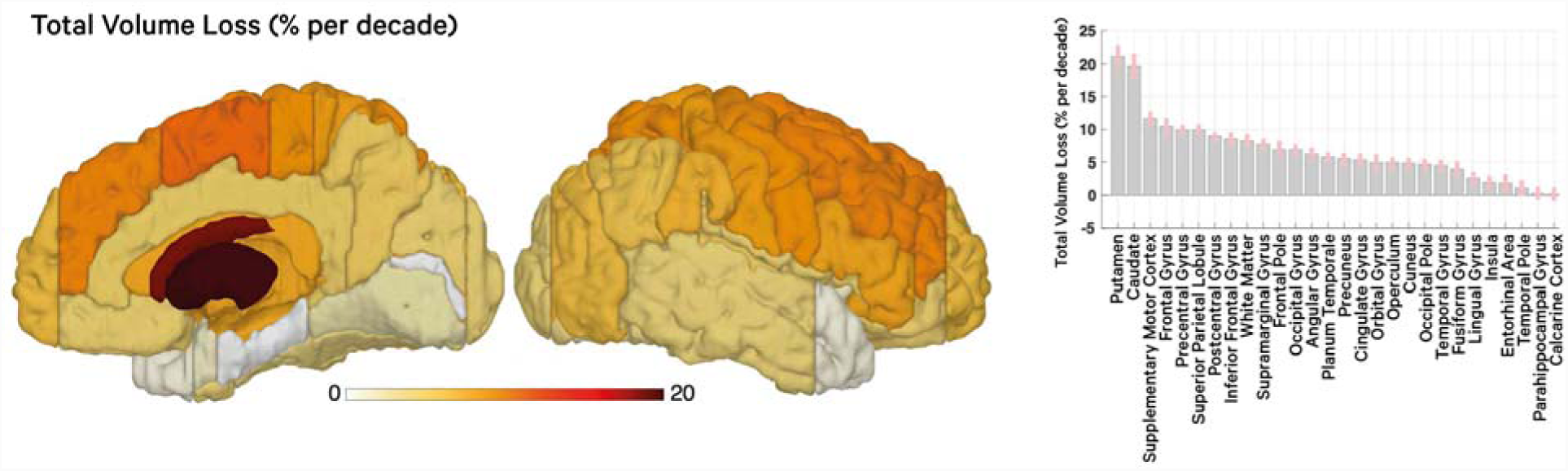
Total brain atrophy during Huntington’s disease motor conversion. Surface projection (left panel) and parameter plot ± SD (right panel) of the overall percent volume loss of regional brain tissue per decade around diagnosis as predicted by the disease progression model estimated using 286 longitudinal MRI scans from 49 HD patients (illustrated in Supplementary Figure 2A). Group level predictions for total volume loss (in % per decade) are shown for the dynamic model with the highest evidence (Supplementary Figure 2B) accounting for individual variations due to age, sex, CAG and confounds (see methods and supplementary information).

### Progression of atrophy shows periods of acceleration within the sensory-motor cortex

Next, the temporal evolution of volumes was studied to differentiate between regions showing a constant rate of volume loss and those exhibiting accelerations or decelerations of shrinkage. The largest rates of constant (linear over time) atrophy were seen in the striatum. Within the cortex, frontal and motor regions showed the greatest rate of volume reduction (Figure 2A).

**Figure 2:**
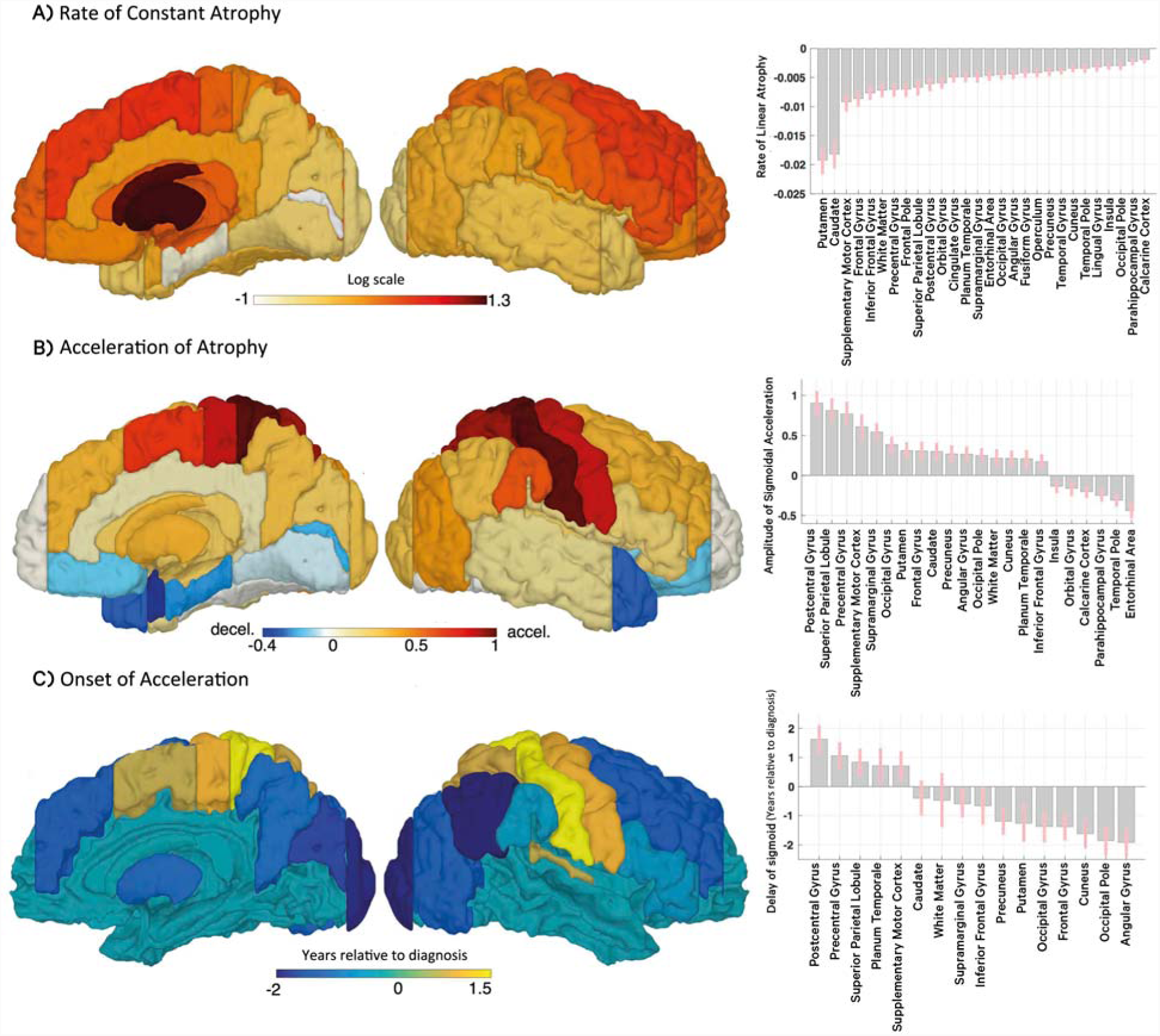
Rates of atrophy and their changes during Huntington’s disease motor conversion. **(A)** Rate of atrophy (or decay rate) of a region indicating constant tissue loss over a decade around HD motor onset. Decay rate refers to parameters of self-connections of regional volume states (Supplementary Figure 2A, see methods and supplementary information). Left panel uses log scale for illustration. **(B)** Regions showing significant accelerations (red) or decelerations (blue) of atrophy above and beyond constant rates of atrophy. Model comparison revealed that highest evidence for model with sigmoidal inputs regional causing accelerations/decelerations (Supplementary Figure 2B & C). Shown is the regional amplitude parameter of the input (for details see Supplementary Figure 5). **(C)** Onset of acceleration (in years relative to diagnosis) of regions that show accelerations of atrophy. Shown is the regional time-shift parameter of the sigmoidal input (for details see Supplementary Figure 5). All results are group level estimates based on longitudinal dynamic modeling and account for effects of age, sex, CAG and confounds.

There were also periods of accelerated volume loss within some regions of the cortex. Accelerations were particularly evident in posterior sensory-motor regions, including the postcentral gyrus, the superior parietal lobule, the precentral gyrus and the supplementary motor cortex (Figure 2B). Minor accelerations of atrophy were also seen in frontal and occipital regions and within the caudate and putamen. In contrast, minor deceleration, i.e. a slowing down of tissue loss, was observed in temporal and some medial-occipital regions. All disease progression model trajectories are summarized in Supplementary Figure 3 (see Supplementary Figures 4 & 5 for data plots and details on models).

We also aimed to address *when* the atrophy progression accelerates with respect to the year of diagnosis. Our analysis suggested that the *timing* of progression also varied across the cortical mantle and subcortical regions (Figure 2C and Supplementary Figures 3 & 5). Taken together, putamen and especially occipital regions showed acceleration of atrophy *prior* to disease onset while sensory-motor cortex showed more emphasized acceleration shortly *after* motor diagnosis.

**Figure 3.**
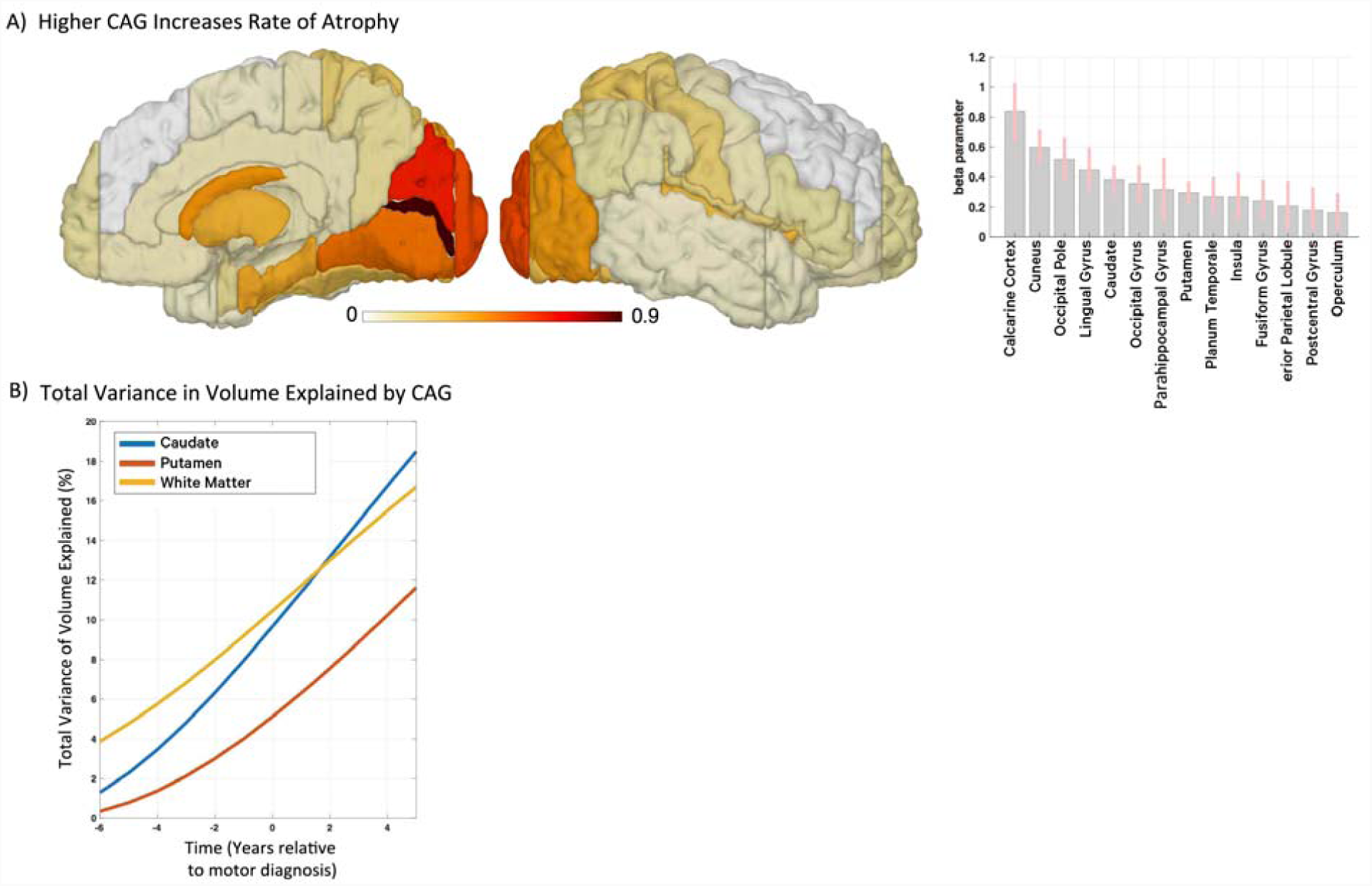
CAG repeat length is related to rate of cortical and striatal atrophy. **(A)** A brain surface projection (left) of parameter ± SD (plot in right panel) that indicates whether individual CAG repeat length predicts regional rate of atrophy/decay. Participants with higher CAG repeat length show increased rate of atrophy (or decay), especially in posterior cortical and striatal areas. Analysis from second level model accounting for effects of age, sex and confounds (see methods and supplementary information). **(B)** The effect of CAG was found to explain an increasing amount of variance of total volume differences across subjects in the caudate, putamen and white matter. Since no comparable increase was observed in other cortical areas this finding is suggestive of individual volume trajectories following more deterministic patterns in striatal and white matter areas. X-axis: disease progression time in years relative to individual motor diagnosis.

### Brain atrophy is related to genetic burden in many regions

We further analyzed the link between CAG repeat length and atrophy across the measured regions. Individual CAG repeat length predicted the rate of atrophy in striatal and occipito-temporo-parietal cortex regions (Figure 3A), suggesting a greater vulnerability of the occipital lobe to increased genetic burden than other regions. Moreover, systematic effects of CAG repeat length on progression were reflected in an increasing amount of variance of atrophy explained by gene differences (Figure 3B).

### No evidence for Inter-regional progression

To explore potential disease spread within the cortex over the 10 year period, we compared models that enable associations of atrophy dynamics between subcortical-cortical and cortical-cortical regions. More specifically, we included between-region connections in the analysis to test whether the atrophy state in one area (e.g. occipital lobe) is predictive of volume change in another (connected) brain area (e.g. sensory-motor cortex). However, model comparisons revealed highest evidence for models without inter-regional interactions (Supplemental Figure 6 and methods section), suggesting that either the regional pattern of atrophy is better described independently, i.e. without significant interactions happening within the morphometric domain, or that the spread of atrophy during HD progression follows a more complex pattern.

### Cortical atrophy can be tied to individual symptom changes

Finally, to evaluate specifics of how regional brain atrophy might contribute to emerging motor symptoms, we extended the HD progression model to a longitudinal brain-behavioral framework (for model information, see Supplementary Figure 7, methods and supplementary information) including both (1) brain volumes (Figure 4B left panel); and (2) motor assessments using TMS scores (Figure 4B right panel). Interestingly, atrophy of the supplementary motor cortex and the caudate was found to be especially predictive of individual motor deficit worsening during progression towards HD (Figure 4A & B and Supplementary Figure 8). The findings align with the notion that neuronal tissue loss within fronto-striatal motor networks might be contributing to worsening motor symptoms. This provides longitudinal evidence for a brain-behavior coupling quantified via an independent motor task but also establishes face-validity of our results.

**Figure 4.**
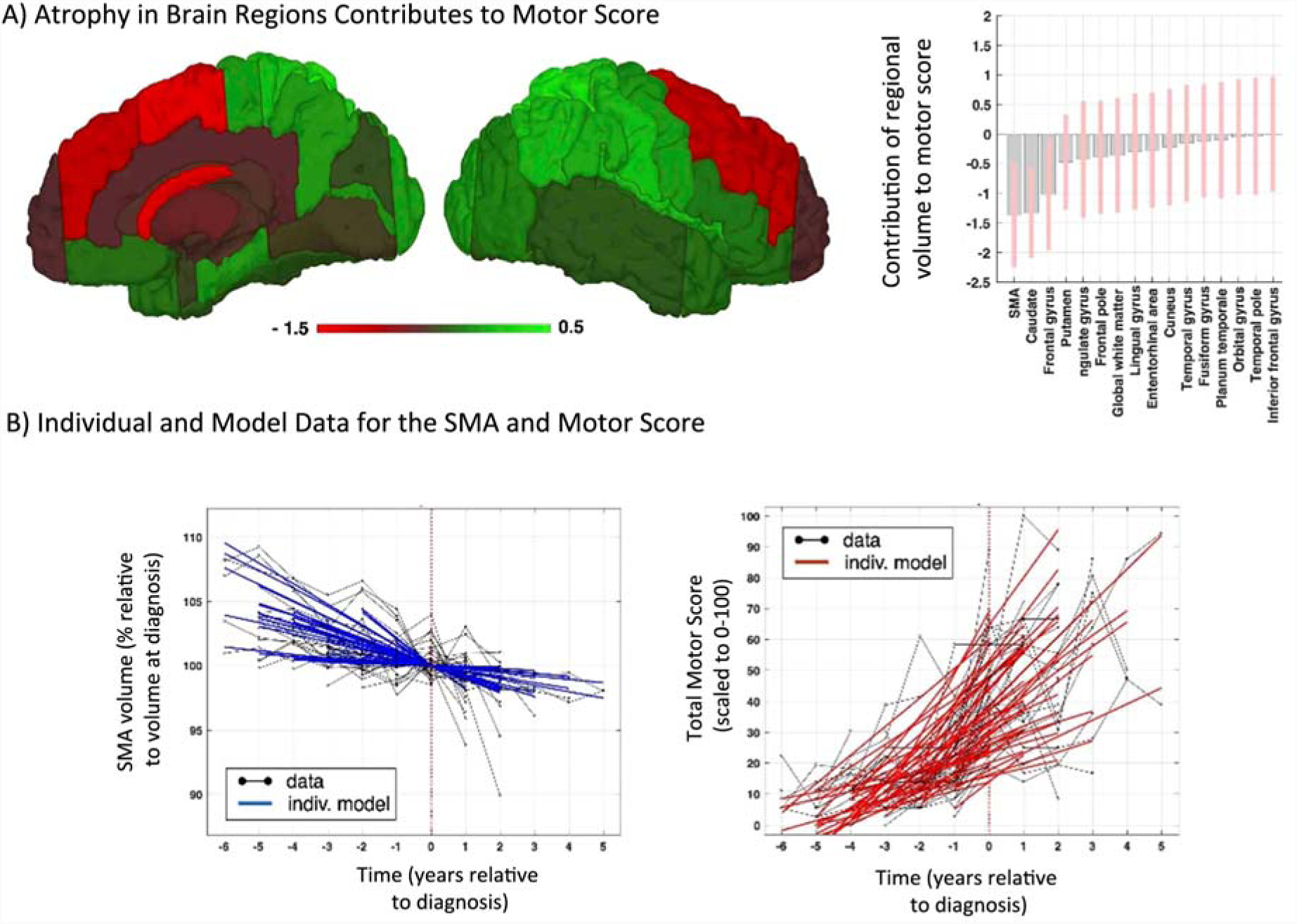
Brain-behavioral model predicts symptom changes during transition to HD. **(A)** Surface projection of weights (left panel) and plot (right panel) that indicate whether a brain region contributes to the prediction of longitudinal motor scores over all timepoints (for model illustration and details see Supplementary Figure 7). Results from second (group-) level accounting for effects of age, sex, CAG and confounds (see methods and supplementary information). **(B)** The left panel shows Supplementary Motor Cortex (SMA) atrophy data (grey, % volume relative to diagnosis) and individual model predictions (blue); the right panel illustrates TMS scores (scaled to 0-100) and individual predictions of our HD progression model. X-axis: disease progression time in years relative to individual motor diagnosis.

## Discussion

Huntington’s Disease is a devastating neurological condition with a complex interplay of physiological, neuronal and behavioral changes^6^. This study provides the first detailed characterization and effect sizes of long-term cortical atrophy over a critical period in HD progression-the onset of clinical motor symptoms. Using a novel dynamic anatomical modeling framework in a unique multi-site longitudinal sample, we explore group level volume trajectories and their individual variability within a broad set of cortical brain regions, covering changes over a ten-year period.

Our findings suggest that volumetric changes are highly variable across the cortical mantle both spatially and temporally. During the period of HD motor symptoms approaching clinical diagnostic levels, frontal and motor areas exhibited the most pronounced total atrophy with rates of >1% volume loss per year in the supplementary motor cortex and dorsal frontal areas; in contrast to comparably preserved anterior and medial temporal lobe regions. This is line with previous studies suggesting that pathological changes in HD are characterized by region-specific vulnerabilities^7,8,11,20^. Mutant HTT is thought to result in neuronal dysfunction via a number of routes; including its effects on cellular proteostasis, transcription, translation and mitochondrial function, as well as via the formation of aggregates^21^. However, these mechanisms and the mechanisms that cause neural atrophy are not well understood. It has been demonstrated that early degeneration occurs in cortico-striatal tracts that are the most highly connected and experiencing the most neural traffic, which are also tracts connecting the basal ganglia to regions within the motor and association networks^22,23^. In the current study the cortical regions that showed the greatest atrophy were the frontal and parietal regions, which have involvement in these networks, offering further evidence in support of the WM findings.

Despite the evidence of widespread atrophy over the course of HD, the widely recognized heterogeneity in the presentation of cognitive and psychiatric disturbances that begin to appear many years prior to motor symptoms^24^ suggests variability in the temporal patterns of regional atrophy. Here, by quantitatively exploring non-linear dynamics during long-term HD progression, we could test for potential accelerations and decelerations in the rate of cortical volume loss that may relate to the variable onset of symptoms. We found that cortical sensory-motor network regions exhibit a significant increase of the rate of atrophy, i.e. an acceleration of shrinkage, occurring shortly *after* motor diagnosis. This is particularly striking and calls into question how cortical changes might relate to individual HD symptomatology. White matter degeneration in pathways between subcortical and motor regions during pre-HD might lead to development and onset of motor symptoms, in turn resulting in an increased rate of cortical volume loss in motor cortical regions, as observed in our study. Increased cognitive and psychiatric symptoms during an earlier premanifest stage, may be related to a period of accelerated frontal lobe atrophy prior to the one studied here which becomes steadier (more constant) around the point of clinical diagnosis. This would require mapping change within participants further from onset.

While primarily interested in cortical progression, we also explored striatal change, which is central to HD pathology. We observed the greatest volume reduction in striatal regions, with both the caudate and putamen showing total loss of around 20% over the ten-year period. There was only minor indication of accelerations in striatal loss. Given the robust evidence showing that subcortical regions undergo atrophy beginning long before motor onset^25^, our findings support previous evidence and thus the validity of the dynamic modeling framework in uncovering the spatio-temporal pattern of volume loss in neurodegeneration.

A longitudinal approach not only allows inference of group level changes but also the contribution of risk factors to apparent individual differences of progression. CAG repeat-length is the key marker of genetic burden in HD and significantly impacts the time of individual clinical disease onset^6^. We found that CAG-length was most predictive of individual rate of volume decline within occipital regions; supported by previous studies demonstrating a link between higher CAG repeat length and HD progression^17,26^ particularly within subcortical and occipital regions^13,27,28^. Interestingly, we also found that some occipital regions, which typically demonstrate substantial atrophy in both pre-HD and manifest-HD^7,9,13,29^ showed relatively low levels of cumulative atrophy. However, previous analyses of occipital atrophy have failed to control for CAG repeat length^7,10^ and longitudinal morphometry may not be optimized for highly myelinated areas^30^. This association between CAG repeat length and occipital atrophy is suggestive of a biological mechanism whereby early visual regions are impacted by genetic burden more than other brain regions.

Using our dynamic modeling framework, we extensively explored the hypothesis of between-region ‘spreading’ of atrophy leading to spatio-temporal patterns of successive impairment. We tested for interactions of change between subcortical and then cortical regions, and/or across between cortical regions during HD progression. Interestingly, there was no detectable evidence of any inter-regional patterns of progression. It is likely that inter-regional interactions follow more complex processes involving subcortical, WM and cortical areas, plus possible time-lagged effects of degeneration; this could be evaluated by including multi-modal and micro-structural states within our modeling framework in future.

Finally, our model temporally relates latent neural (state-) changes to behavioral changes, rather than simply correlating cortical atrophy with behavioral symptoms across patients^8,31^. By linking changes in structure and behavior, we can associate pathological change with patients’ experience of symptom progression. Motor network atrophy, for example, is both associated with clinical motor onset and also acts as an independent predictor of motor performance. We found that atrophy within the supplementary motor area (SMA), frontal gyrus and caudate contributed most to worsening performance on a motor task. Both the SMA and frontal lobe are involved in movement planning and are highly integrated with the basal ganglia^32^, signifying that these regions are particularly relevant for HD progression and motor symptom onset. Furthermore, the association between cortical atrophy and motor score demonstrates face validity of our novel modeling technique for quantifying neurodegeneration.

The modeling framework we used here was developed to address weaknesses in previous analysis methods and approach the quantification of GM change via a dynamic systems method^5^ using a unique longitudinal HD cohort. The ability to quantify both constancy and changes in the rate (accelerations/decelerations) of cortical GM atrophy over time, while accounting for individual variability over multiple timepoints offers a more powerful approach to structural MRI modeling than many previous methods^33,34^. It is important to consider that the use of motor diagnosis for temporal alignment of all participant’s progression in our model introduces a level of subjectivity as to inclusion of participants, plus focus on participants within 6 years to motor onset prevents capture of earliest cortical atrophy patterns. However, we carefully account for differences due to age, sex, TIV and scanning site to render the group inference unbiased. Nevertheless, the regional progression observed in these results during a particularly crucial period in HD progression is clinically meaningful, supported by post-mortem studies, previous imaging data and disease symptoms.

Our findings provide the most detailed characterization of cortical atrophy in neurodegenerative disease within an HD cohort at the point of clinical diagnosis. By applying a recently validated model that is uniquely able to map temporal and spatial cortical change, we have demonstrated that cortical atrophy shows regional variability related to genetic factors and predicts worsening performance on a motor task, the key HD phenotype. Brain regions that appear to undergo the greatest atrophy during this period are typically associated with clinically reported symptoms, with prominent accelerations in atrophy within the motor cortex suggesting a close relationship between symptom onset and cortical change. This work represents a principled approach to modeling longitudinal structural MRI data that offers new insights into the spatial and temporal phenotype of cortical changes, and in turn the biological underpinnings of neurodegeneration.

## Methods

### Participants

Participants were from both the longitudinal multi-site observational TRACK-HD and TrackOn-HD study cohorts^7,18^. For TRACK-HD, participants attended four yearly visits and were divided into control, Pre-HD and manifest HD groups at recruitment^7^. TrackOn-HD followed on from the TRACK-HD study and included both a subset of TRACK-HD participants and newly recruited participants. Participants who were initially included in the Pre-HD group in either cohort and subsequently transitioned to manifest HD (’converters’) during the course of data collection were included in the current study. These criteria were used to create a group of participants experiencing a similar stage of disease progression.

For TRACK-HD, gene-positive participants were required to have a positive genetic test of ≥40 CAG repeats and a burden of pathology score > 250^35^. Pre-HD participants all had a Total Motor Score (TMS) on the Unified Huntington’s Disease Rating Scale (UHDRS) of < 5^36^, indicating a lack of motor symptoms. Participants were classed as converters if they were recruited into the study as Pre-HD based on the above criteria, and then received a Diagnostic Confidence Score (DCS) score of ≥4 at any subsequent timepoint, indicating that they had met clinical diagnostic criteria for manifest HD. Fifty participants met this criterion for conversion during TRACK-HD/TrackOn-HD. One additional participant received a DCS of 4 at TRACK-HD visit 4, but then reverted to DCS <4 at a later timepoint and was excluded from this investigation.

Data for all converters were re-aligned to consolidate year of motor conversion across all participants (Supplementary Figure 1). This was done to increase homogeneity of disease progression within the group and to define a comparable disease progression time variable. The first year of DCS = 4 was designated as year of conversion (timepoint = 0), and each year prior to conversion labeled as year-1,-2,-3, etc. Every year after conversion was labeled as year 1, 2, 3, etc. Individual variability of changes beyond the synchronizing event of motor diagnosis will be accounted for during modeling (see below and supplementary information).

Total Motor Score (TMS) was used to approximate clinical motor progression^36^. TMS is part of the Unified Huntington’s Disease Rating Scale (UHDRS) and is a clinical rating scale performed by a physician. TMS measures the presence of motor symptoms, and ranges from 0-60 with a score of <5 indicating no substantial motor symptoms. The TMS rating scale requires participants to perform various motor tasks (for example, tandem walking) while being observed by a trained clinician, and shows high reliability^36^.

### MRI data acquisition

3T T1-weighted scans were acquired from four scanners for both TRACK-HD and TrackOn-HD, and acquisition protocols were the same for both studies. Two scanners were Siemens and two were Philips. The parameters for Siemens were TR = 2200ms, TE = 2.2ms FOV = 28cm, matrix size = 256×256, 208. For Philips TR = 7.7ms, TE = 3.5ms, FOV = 24cm, matrix size = 242×224, 164. The acquisition was sagittal to cover the whole-brain. There was a slice thickness of 1mm, with no gap between slices. These acquisition protocols were validated for multi-site use^7^. Scanners were monitored over time to ensure consistent acquisition of images and all images were visually assessed for quality at time of data collection; specifically artifacts such as motion, distortion and poor tissue contrast (IXICO Ltd. and TRACK-HD imaging team, London, UK).

### Longitudinal image processing

An initial longitudinal within-participant registration pipeline from SPM12 was used to create an average image for each participant^37^. This process included between-timepoint scan registration, creation of an average participant scan for all timepoints and differential bias correction for between-timepoint scan inhomogeneity^38^. Registration was performed using default settings and visual quality control (QC) was performed on all registered scans. Each average scan was then parcellated into 138 regions using MALP-EM, a fully automated segmentation tool^39^ validated for use in HD^19^; previously manually segmented whole-brain regions were binarized and included in the MALP-EM pipeline to improve brain extraction^40^. Each average segmented region was then multiplied by Jacobian deformation maps (derived from the registration step) to create a volumetric map for each region for each timepoint. All segmentations underwent visual QC. One dataset did not pass QC due to errors in segmentation that could not be rectified.

The 138 regions were combined into 55 larger regions based on spatial localization and visual inspection to reduce noise within small cortical regions (Supplementary Table 1). 50 cortical regions (25 bilateral pairs), four subcortical regions (bilateral caudate and putamen) and one global white matter volume were included in the analysis. To facilitate clear across-region comparisons, regional brain volume changes relative to volume at timepoint of motor diagnosis (in percent) were analyzed.

### Hierarchical disease progression models

The previously established framework for dynamic modeling of longitudinal structural MRI was applied to regional volume data to describe neuronal tissue loss over progression time in HD^5^ using an empirical Bayes framework^4^. We briefly describe the methods here; see supplementary information for further details.

The applied disease progression model is illustrated in Supplementary Figure 2A. Model development involved Bayesian model comparisons of seven models of cortical atrophy to find which best described the progression of cortical atrophy in our HD cohort. Across the seven models both linear and non-linear (acceleration or deceleration) patterns of atrophy with varying system inputs (linear, piecewise linear, sigmoidal) were explored (Supplementary Figure 2B). Each of these models were estimated at the first-level for each individual and inverted using Variational Laplace methods^16^. The parameter estimation (or system integration) modeled measures of volume in each brain region symmetrically from point of clinical diagnosis to the earliest timepoint prior to diagnosis and the latest timepoint after motor diagnosis.

Hierarchical modeling increases power for detecting group-level effects by modeling differences in first-level parameters and their uncertainty, and accounting for differing number of visits. Therefore, the seven models for each participant were embedded in a larger second-level model to estimate group-wise change. Full hierarchical (individual and group level) models for each of the seven patterns of atrophy were estimated, incorporating both the second level group mean and covariates including CAG repeat length, sex, age (at motor diagnosis), TIV and scanning site (age was orthogonalized with respect to CAG, due to the high correlation between the two measures). Bayesian Model Selection (BMS) was then applied to select which of the seven models or patterns of atrophy best explained our observed data across the group. BMS both optimizes model fit while penalizing complexity and is, therefore, appropriate for use in highly parameterized hierarchical disease progression models^41^. Of the seven models examined, model evidence (measure of the best model fit) was highest for a dynamic model (with state equation *dx/dt=Ax+Cu)* using regional sigmoidal inputs (*u*) allowing a specific delay and amplitude of acceleration/decelerations (Supplementary Figure 2B for illustration of different inputs, Supplementary Figure 2C for model evidence, and Supplementary Figure 4 for estimated sigmoidal inputs obtained in the winning model). Results for this model are subsequently reported in terms of the group-derived posterior distribution (mean±SD).

The generative disease progression model (illustrated in Supplementary Figure 2A, with results presented in Figures 1-3) was restricted to describe volumes in multiple brain regions only. Notably, the original model did also not allow for inter-regional dynamics i.e. the connection parameters were fixed to zero. In order to test for potential inter-regional dynamics of regional morphometry during HD progression we explored various striatal-cortical and cortical-cortical networks (details and illustration in Supplementary Figure 6) using weakly informative priors on connection weights. However, highest model evidence was observed for uncoupled models and thus all main findings were restricted to models with region-specific self-connections (diagonal of *A* matrix) and contribution of inputs (C matrix). Finally, we extended the observational model to predict longitudinal TMS rating using a regression approach based on linear combination of all dynamic brain state variables (for more details see Supplementary Figure 7 and supplementary notes on modeling).

### Code availability

The link to custom made scripts and synthetic example dataset that demonstrate SPM-based dynamic modeling of longitudinal HD data applied during this study can be provided upon request to the corresponding authors. Please open readme.txt for further details on how to apply the code. Please note that this particular code has not been thoroughly tested with other software versions than MATLAB2016b, SPM12 r7355 and other parameter choices might produce errors during processing attempts. The code aims at transparency and illustration but is not intended for clinical use. It is free but copyright software, distributed under the terms of the GNU General Public License as published by the Free Software Foundation (either version 2, or at your option, any later version). Further details on “copyleft” can be found at http://www.gnu.org/copyleft/. In particular, software is supplied as is. No formal support or maintenance is provided or implied. For any questions and requests please contact

## Author contributions

TRACK-HD Investigators designed the experiment. E.J., G.Z., S.G., R.S. and W.P. conducted the experiment and analyzed the data. E.J., G.Z., and S.G. wrote the paper. G.R. edited the manuscript. S.T. was Principal Investigator of the TRACK-HD and TrackOn-HD studies, and edited the manuscript.

### Acknowledgments

The authors wish to thank the TRACK-HD and TrackOn-HD study participants and the CHDI Foundation, a not-for-profit organization dedicating to finding treatments for HD. Some of this work was undertaken at University College London Hospital/University College London, which received funding from the Department of Health NIHR Biomedical Research Centres funding scheme. EBJ, SJT, RIS and SG are supported by a Wellcome Trust Collaborative Award (grant code 200181/Z/15/Z). GZ would like to thank Emrah Diizel for support at the DZNE.

## TRACK-HD Investigators

AUSTRALIA Monash University, Victoria: SC Andrews, JC Campbell, M Campbell, E Frajman A O’Regan, I Labuschagne, C Milchman, C Pourchot, S Queller, JC Stout; CANADA University of British Columbia, Vancouver: A Coleman, R Dar Santos, J Decolongon, B Leavitt, A Sturrock FRANCE APHP, Hopital Salpetriere, Paris: E Bardinet, A Durr, C Jauffret, D Justo, S Lehericy, C Marelli, K Nigaud, R Valabrègue; GERMANY University of Münster, Münster: N Bechtel, S Bohlen, R Reilmann; University of Bochum, Bochum: A Hoffman, P Kraus; University of Ulm, Ulm: GB Landwehrmeyer. NETHERLANDS Leiden University Medical Centre, Leiden: SJA van den Bogaard, EM Dumas, J van der Grond, EP t’Hart, C Jurgens, RAC Roos, M-N Witjes-Ane. U.K. St Mary’s Hospital, Manchester: N Arran, J Callaghan, D Craufurd, C Stopford; London School of Hygiene and Tropical Medicine, London: C Frost, R Jones; University College London, London: H Crawford, NC Fox, C Gibbard, NZ Hobbs, N Lahiri, I Malone, R Ordidge, G Owen, A Patel, T Pepple, J Read, MJ Say, E Wild, D Whitehead; Imperial College London, London: S Keenan; IXICO, London: D M Cash; University of Oxford, Oxford: C Berna, S Hicks, C Kennard. U.S.A. University of Iowa, Iowa City, IA: T Acharya, E Axelson, H Johnson, DR Langbehn, C Wang; Massachusetts General Hospital, Harvard, MA: S Lee, W Monaco, HD Rosas; Indiana University, IN: C Campbell, S Queller, K Whitlock; CHDI Foundation, New York, NY: B Borowsky, AI Tobin.

## TrackOn-HD investigators

AUSTRALIA Monash University, Victoria: SC Andrews, I Labuschagne, JC Stout. CANADA University of British Columbia, Vancouver: J Decolongon, M Fan, T Koren, T Petkau, B Leavitt; FRANCE ICM and APHP, Hôpital Salpêtriere, Paris: A Durr, C Jauffret, D Justo, S Lehericy, K Nigaud, R Valabrègue; GERMANY Freiburg University, Freiburg: S Klöppel, L Mincova, Scheller E; George Huntington Institute, Munster: R Reilmann, N Weber; University of Ulm, Ulm: B Landwehrmeyer, I Mayer, M Orth; NETHERLANDS Leiden University Medical Centre, Leiden: SJA van den Bogaard, RAC Roos, EP’t Hart, A Schoonderbeek; U.K. London School of Hygiene and Tropical Medicine, London: A Cassidy, C Frost, R Keogh; Manchester University, Manchester: D Craufurd; University College London, London: C Berna, H Crawford, M Desikan, R Ghosh, D Hensman, EB Johnson, D Mahaleskshmi, I Malone, P McColgan, M Papoutsi, J Read, A Razi, G Owen; U.S.A. University of Iowa, Iowa City, IA: H Johnson, DR Langbehn, Long J, Mills, J. CHDI Foundation, New York, NY: B Borowsky.

## Data Availability Statement

Our participants did not give informed consent for their measures to be made publicly available, and therefore we did not present the underlying data in the manuscript itself. Requests for access to the TRACK-HD and TrackOn-HD data supporting the analyses presented in this paper should be made via the CHDI Foundation. For more information, please contact.

