## Supplementary Information for "Dynamics of cortical degeneration over a decade in Huntington’s Disease"

**­­**

**Content:**

Supplementary Figures 1-8

Supplementary Table 1

Supplementary Notes on Modelling

**Supplementary Figures**

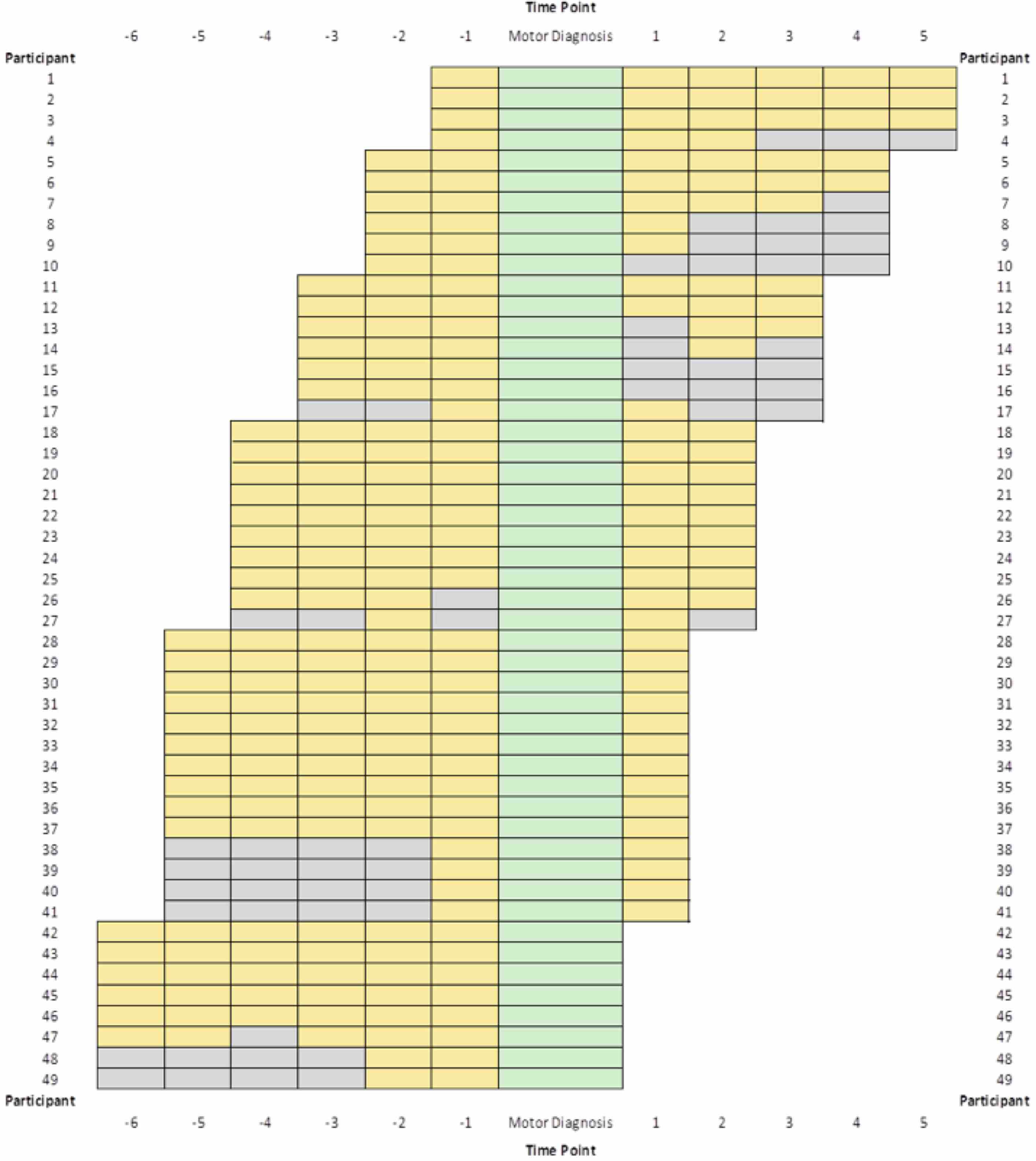

**Supplementary Figure 1**. **A schematic showing all MRI scan acquisitions included in analysis.** Data is shown re-aligned so motor diagnosis is consistent across participants. Green represents motor diagnosis, yellow represents available MRI data and grey represents missing data. Missing data includes time-points for which a participant was not yet recruited (e.g. when a participant was recruited at baseline of TrackOn-HD, such as Participant 49), or had dropped out of the study (e.g. Participant 16 dropped out at the end of TRACK-HD and did not participate in TrackOn-HD), and when a participant could not attend a time-point (e.g. Participant 26). X-axis: disease progression time (in years relative to individual motor diagnosis) also used for dynamic HD modelling.

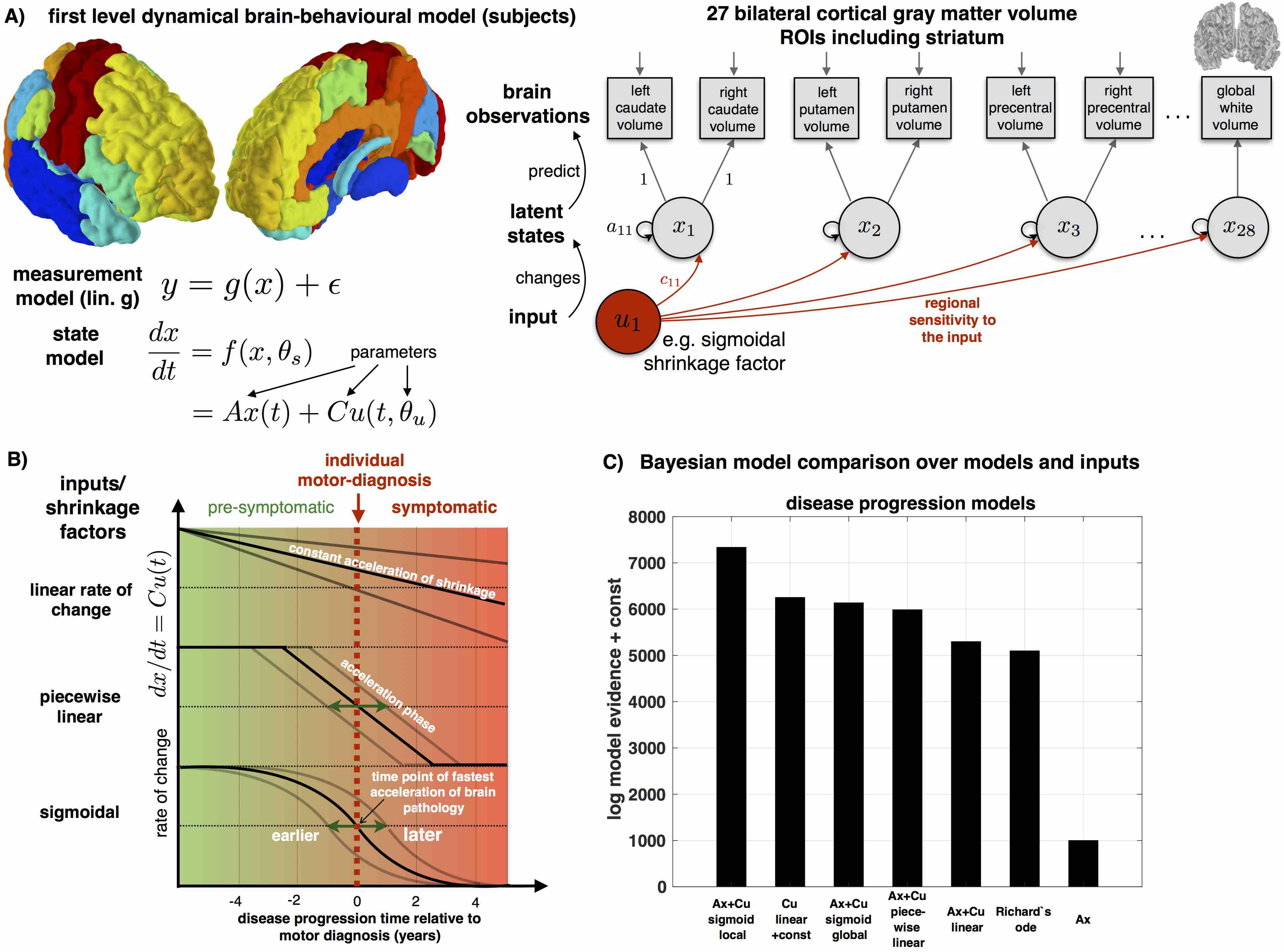

**Supplementary Figure 2. Overview of applied dynamic disease progression modelling framework.** To describe disease related brain changes in transition phase towards HD, we applied a dynamical systems framework previously established in Ziegler et al.^1^. **(A)** An illustration of first (subject) level model using a parcellation of the brain into 27 bilateral grey matter ROI volume including striatum and the global white matter volume. The generative model is defined by a system (with state model and measurement/observation model) describing changes in 28 regional volume states (all grey ROIs and total white) during the period from -6 years before to 5 years after clinical motor diagnosis. **(B)** Illustration of explored system inputs causing different forms of acceleration of pathology during transition from pre-symptomatic to symptomatic disease phase. In case of presence of non-linearities, the rate of change (velocity) of progression might change linearly with progression time (top), only during a restricted period/phase (middle), or transitions smoothly following a sigmoidal shape (bottom, see also Supplementary Figure 5) **(C)** Bayesian model evidence of multiple dynamic models included in the model space for comparison. All models estimated were hierarchical with bottom level for individuals and one top level describing mean group dynamics, covariates and confounds (see Friston et al., 2016 for details on inference using Parametric Empirical Bayes, PEB). We explored models with (and without) several forms of additive input to the system, more complex and also conventional polynomial alternatives. Highest model evidence was found for linear dynamic system having a regional sigmoidal input that allows for additional accelerations/decelerations of volume state changes above and beyond endogenous dynamics (induced by self-connections) (cf. methods and supplementary notes on modelling).

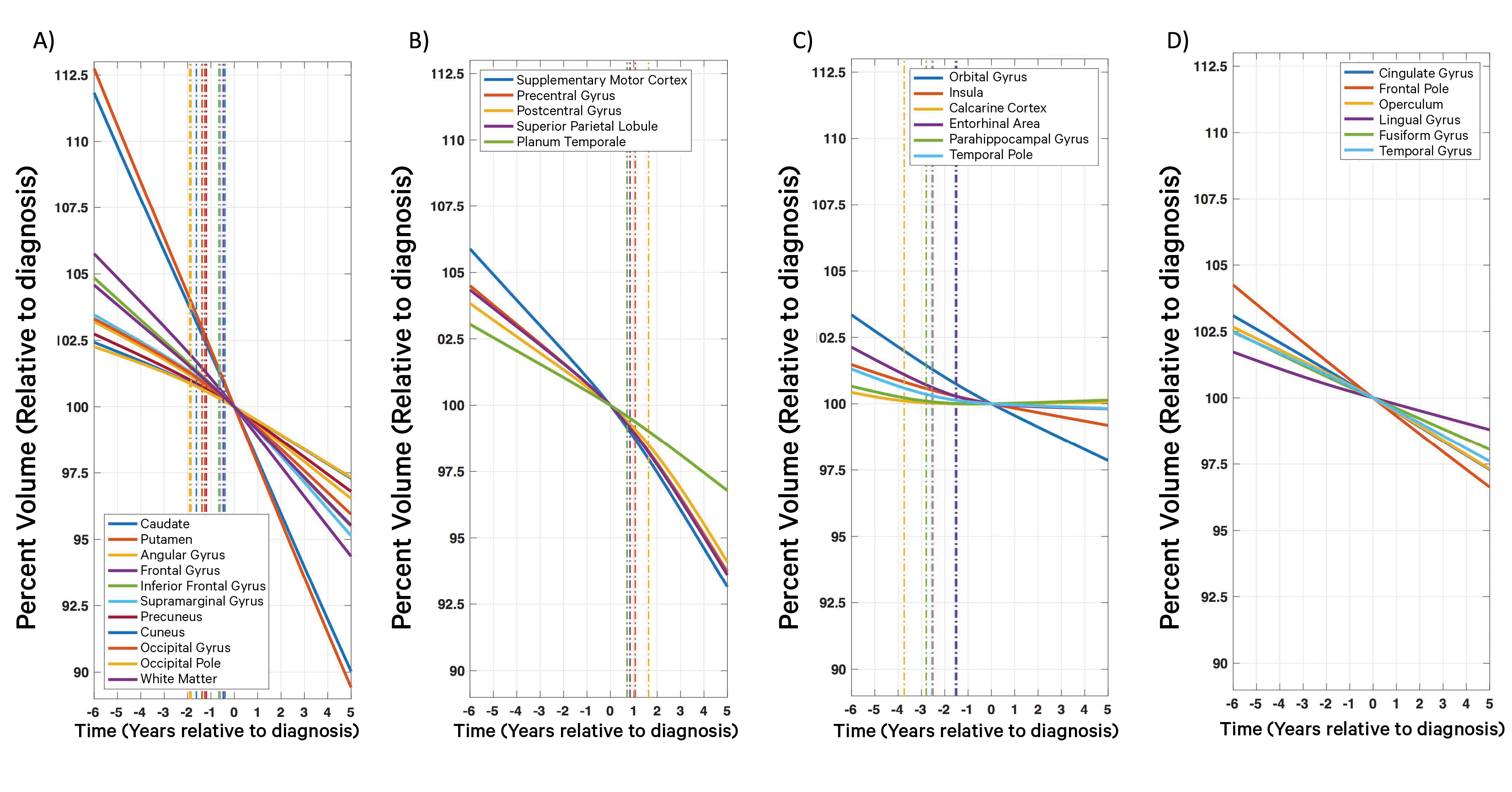

**Supplementary Figure 3. All regional atrophy trajectories over a decade as predicted by the HD progression model.** Regions differ qualitatively with evidence for some ROIs **(A)** showing accelerations before motor diagnosis (i.e. time zero) (see also Figure 2B & C); **(B)** showing accelerations after motor diagnosis; **(C)** showing decelerations; or with **(D)** no indication of non-linear effects. Vertical dotted lines show the estimated time point of strongest accelerations/decelerations of atrophy progression for each region (cf. for data plots see Supplementary Figure 4; cf. more details on accelerations see Supplementary Figure 5 and supplementary notes on modelling). Y-axis: percent volume (relative to volume at time-point of diagnosis). X-axis: disease progression time in years relative to individual motor diagnosis.

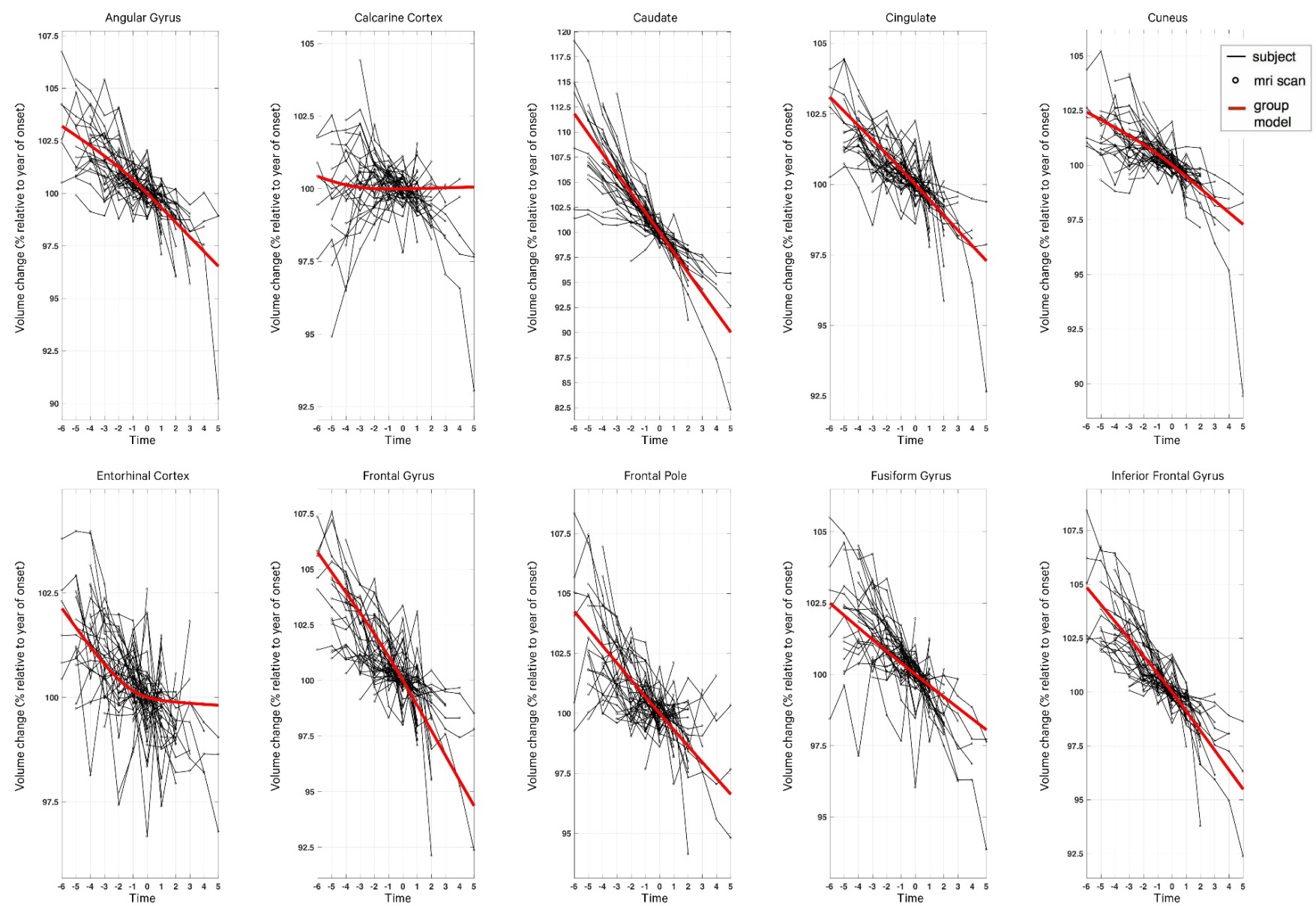

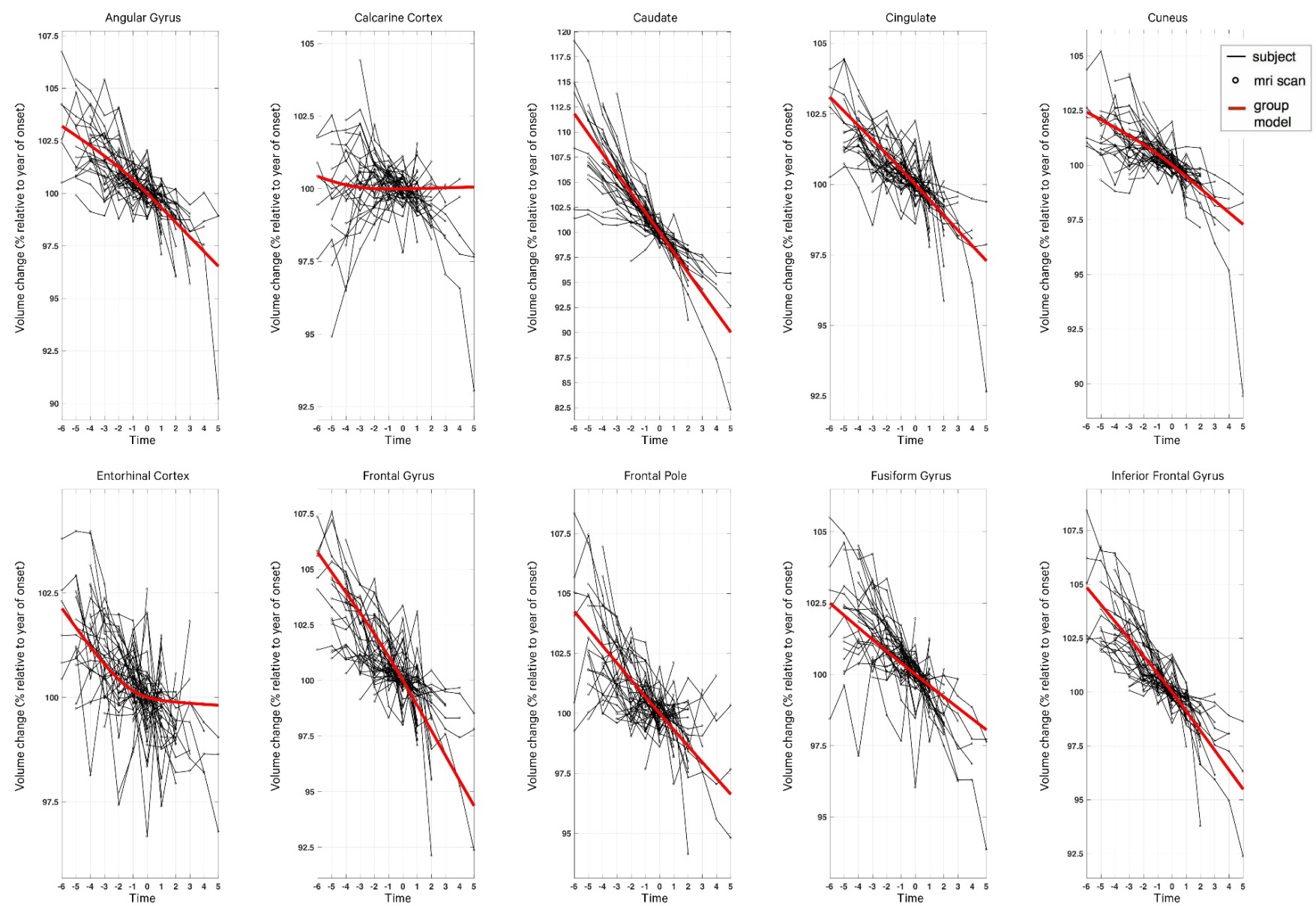

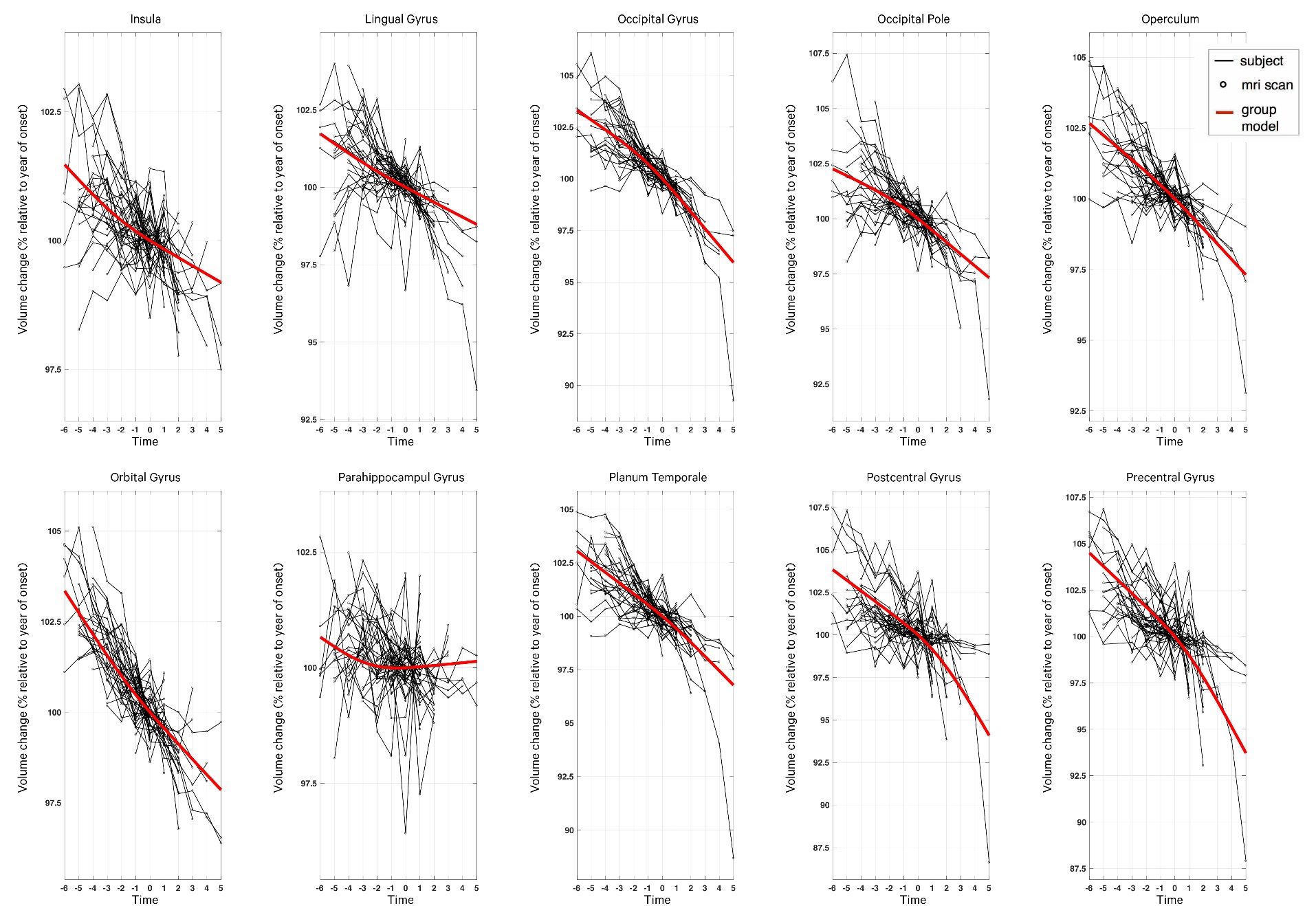

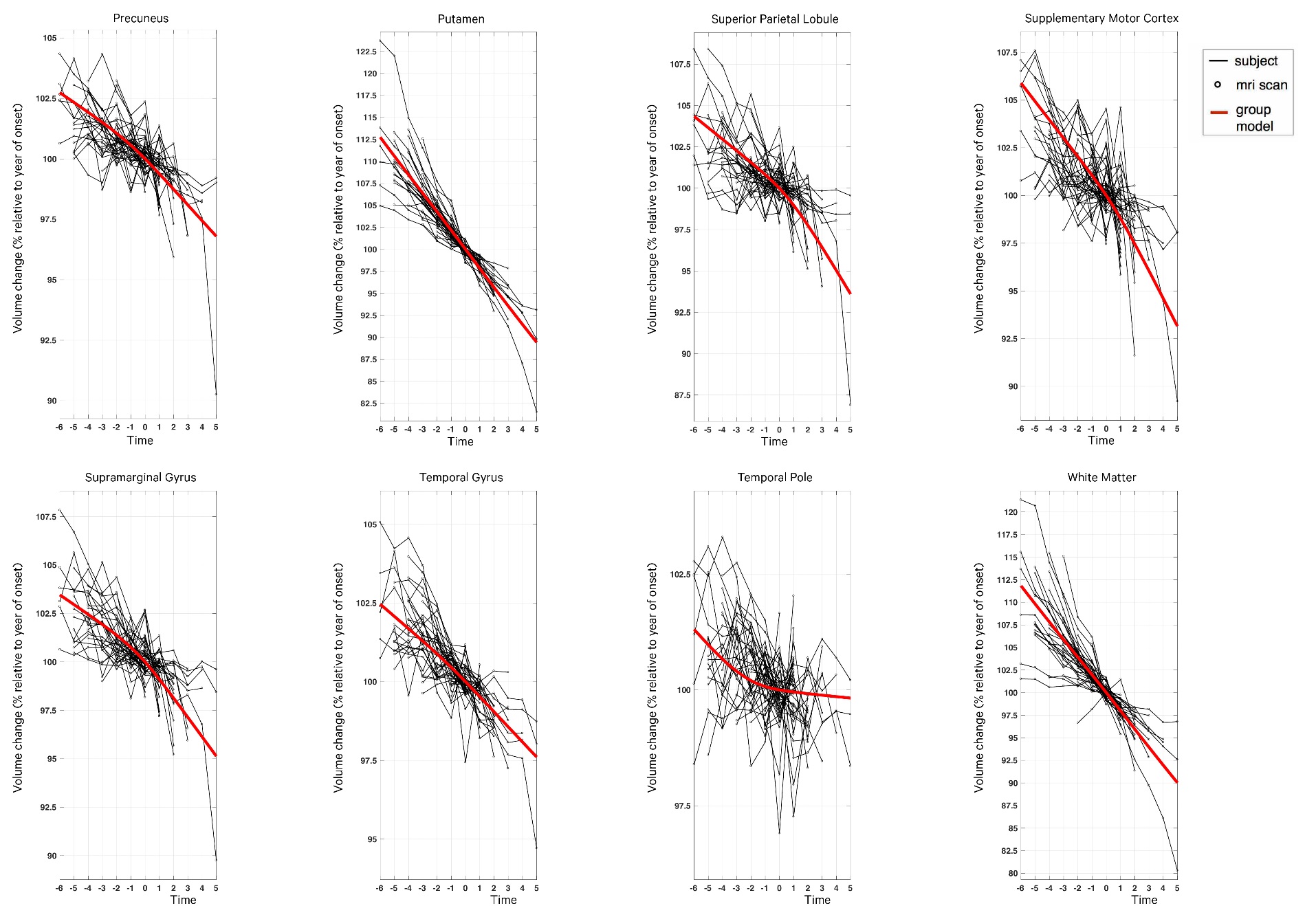
­

**Supplementary Figure 4. Regional volume data from all ROIs and model predictions of the HD progression model.** Plots show regional longitudinal raw data (percent volume relative to volume at year of motor diagnosis) with each thin black line representing one of the 49 participants. The group level predictions from the dynamical HD progression model with highest Bayesian model evidence is shown in red. With exception of the white matter volume, regional volumes refer to the left hemisphere with corresponding right hemispheric volume exhibiting very similar progression (not shown). Y-axis: percent volume (relative to volume at time-point of diagnosis). X-axis: disease progression time in years relative to individual motor diagnosis. For more details on trajectory estimation see methods and supplementary notes on modelling.

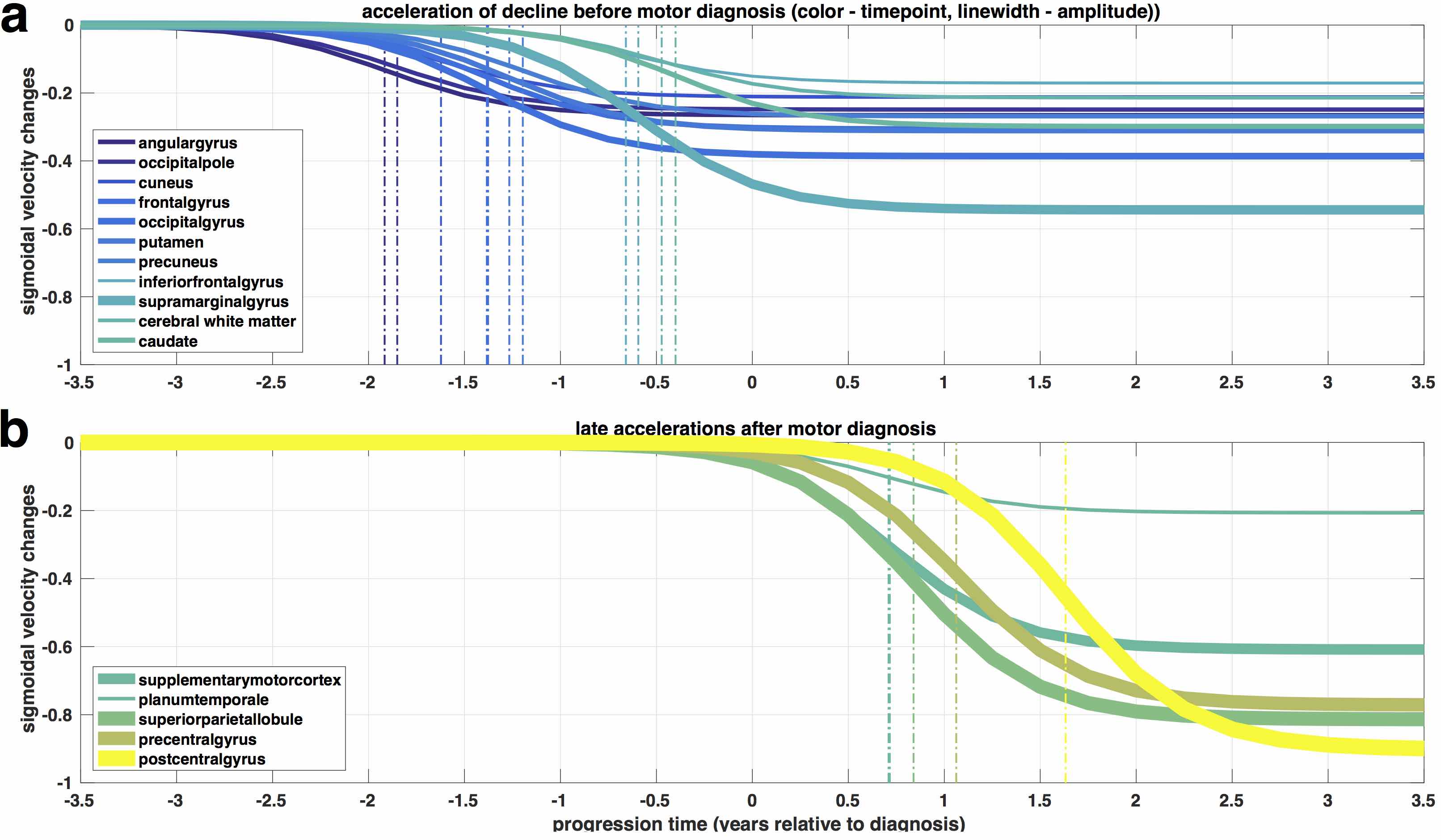

**Supplementary Figure 5. Accelerated disease progression using sigmoids as system inputs.** In addition to approximately linear volume decay implied by self-connections, including a sigmoidal input (cf. schematics in Supplementary Figure 2A & B) resulted in an improvement of Bayesian model evidence (cf. Supplementary Figure 2c for model comparison). This indicated a significant non-linearity of disease progression for some regions. Using the obtained group level parameters, we here illustrate the sigmoidal inputs that contribute to the rate of change (velocity) used during the integration of the dynamical system (cf. methods and supplementary notes on modelling). **(A)** We show sigmoids from brain regions with early accelerations (transitioning from lower to higher rate of shrinkage) which occur *before* motor diagnosis. **(B)** This presents sigmoids from regions accelerating *after* motor diagnosis, e.g. the sensory-motor cortex. Y-axis: For illustration purpose we here only show the contribution of the input to the rate of change but not the contribution of the endogenous self-connection (which causes the constant decay, shown in Figure 2A). Thus, all depicted regions initially progress with rates of change around zero, then accelerate (sooner or later) and end at a slightly higher rate of change (cf. fully integrated volume trajectories are summarized in Supplementary Figure 3 & 4). X-axis: disease progression time in years relative to individual motor diagnosis.

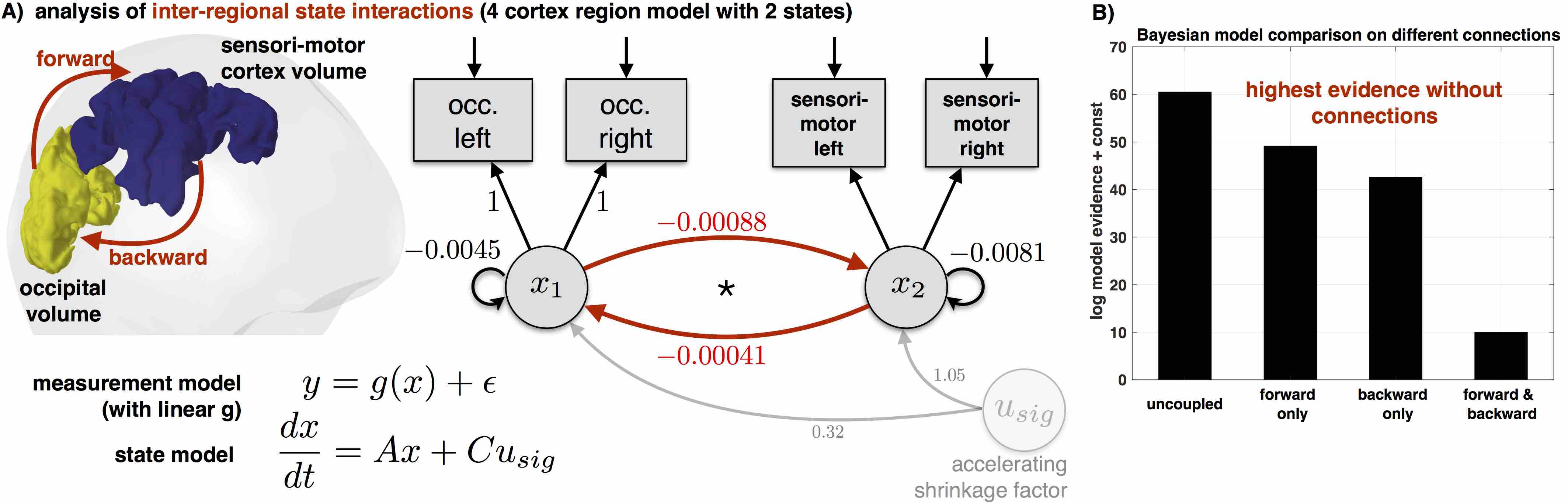
**Supplementary Figure 6. Exploring inter-regional dynamics during HD progression.**
We explored potential disease spreading including regional connections in the dynamical system (state model with connectivity matrix *A,* cf. supplementary notes on modelling). We particularly focussed on striatal-cortical (not shown) and cortical-cortical interactions. **(A)** We hypothesized and tested interactions by focussing on 2 (bilateral) region models with occipital and sensory-motor cortex volume respectively (top left surface projection). Off-diagonal elements of connectivity matrix were used to test whether individual volume state in region 1 (occ.) significantly affects rates of change of progression in region 2 (sensory-motor cortex), above and beyond decay induced by self-connections (i.e. rate of atrophy/decay). Forward, backward and bi-directional connectivities were implemented and included as separate models in the model space. **(B)** Bayesian model comparison revealed highest model evidence for models without additional inter-regional connections. This suggests that spatial and temporal dynamics of volume changes are parsimoniously described using local dynamical parameters and inputs presented in main results (Figure 1-3). Neuronal processes such as micro-structure (not optimally) reflected in macroscopic volumes might be involved in causal unfolding of HD disease pathology (cf. methods and supplementary notes on modelling).

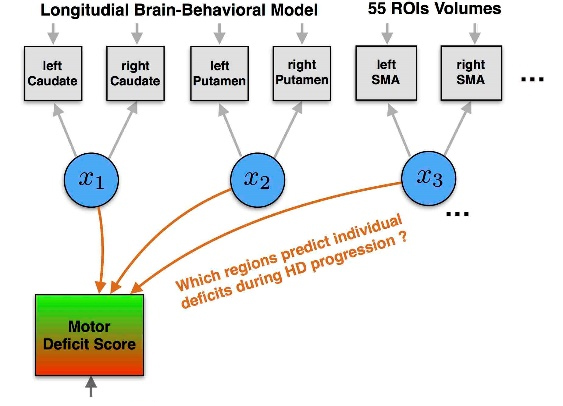

**Supplementary Figure 7. Extending the dynamic model to predict individual symptom changes during transition towards HD.** Illustration of an extended brain-behavioral dynamical model predicting longitudinal observations from two domains, i.e. regional brain volumes (grey boxes) and TMS motor scores (red-green box). The 28 hidden brain states *x(t)* (blue circles) inferred in the generative model (also illustrated in Supplementary Figure 2) were here used to simultaneously predict longitudinal TMS motor scores available for all time-points with MRI scans from 47 of 49 participants. Notably, full model inversion is performed jointly with brain and behavioural data. The TMS prediction is following a multiple linear regression via an extended observational model (linear combination of states with prior weights around zero, cf. methods and supplementary notes on modelling). Larger weights in a region indicate a higher contribution of the (time-varying) regional atrophy to prediction of symptoms during progression (cf. weights shown in Figure 4A). Examples of individual participant model predictions for supplementary motor area (SMA) volume and TMS scores are shown in Figure 4B (cf. caudate and TMS also in Supplementary Figure 8).

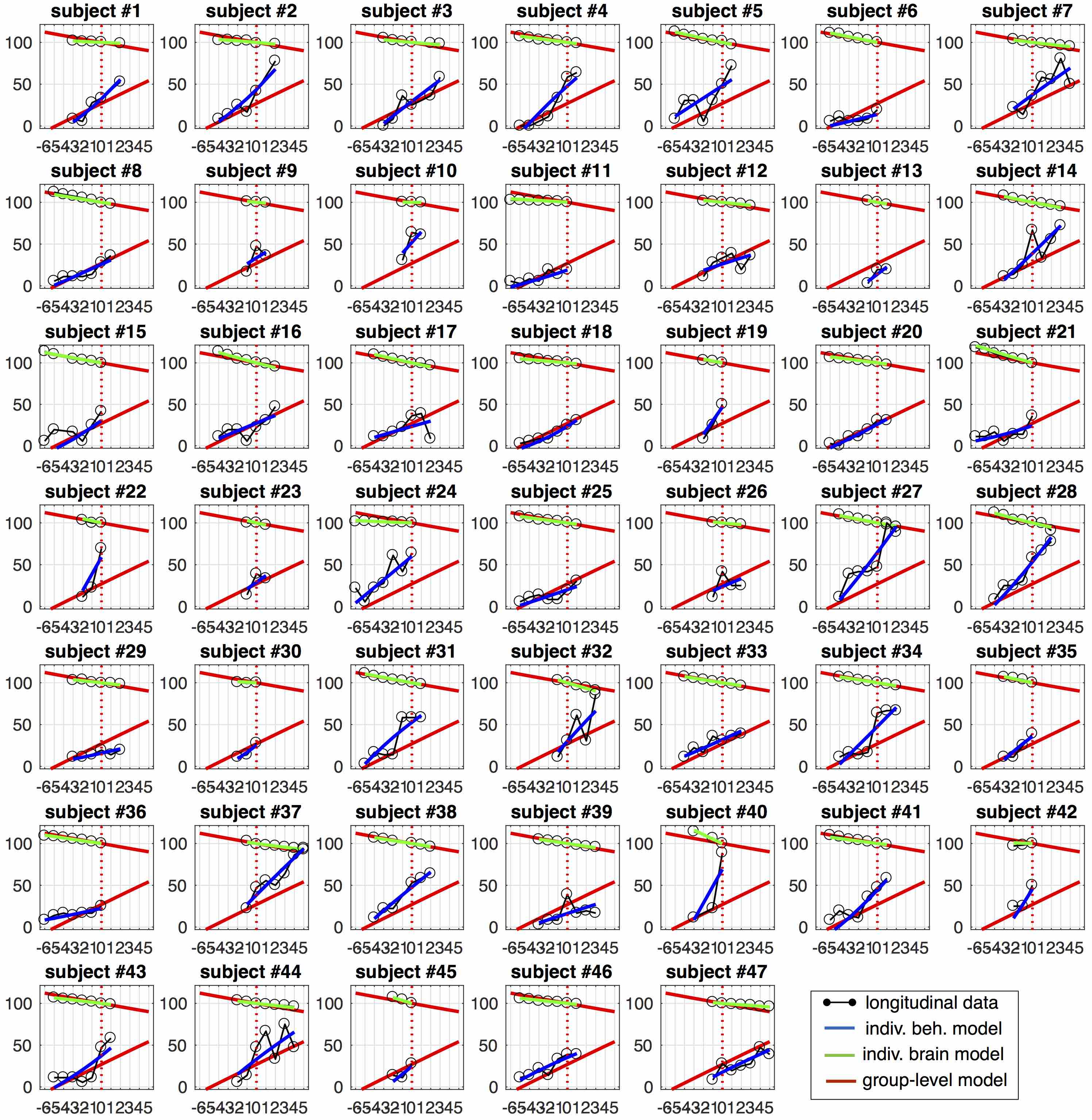
 **Supplementary Figure 8. Example of individual-level model of caudate and motor score disease progression trajectories.** The main generative model predicts individual regional brain volumes during HD disease progression (illustrated in Supplementary Figure 2, cf. obtained parameters shown in Figure 2ABC). The upper part of the plot shows the individual-level (green) as well as group-level (red) model predictions and the observed caudate volumes (black, % relative to motor diagnosis) for 47 of 49 subjects with available MRI scans and motor symptom assessments (TMS). We also extended the model to a generative brain-behavioral model including symptoms (illustrated in Supplementary Figure 7, cf. parameters shown in Figure 4; cf. methods, supplementary notes on modelling). Lower part of the plot shows the TMS observations (black, normalized to [0, 100]) and corresponding individual-level (blue) and group level model predictions (red) of all 47 subjects with available data. Y-double-axis: percent volume (relative to volume at time-point of diagnosis) or TMS scaled to 0-100. X-axis: disease progression time in years relative to individual motor diagnosis.

**Supplementary Table 1.** The final regions measured in this study, and the original regions output by MALP-EM that were combined to create the final regions.

| **Final Region Name** | **Original Combined Regions** |
| --- | --- |
| Angular Gyrus Right | Angular Gyrus Right |
| Angular Gyrus Left | Angular Gyrus Left |
| Calcarine Cortex Right | Calcarine Cortex Right |
| Calcarine Cortex Left | Calcarine Cortex Left |
| Cuneus Right | Cuneus Right |
| Cuneus Left | Cuneus Left |
| Entorhinal Area Right | Entorhinal Area Right |
| Entorhinal Area Left | Entorhinal Area Left |
| Frontal Pole Right Occipital | Frontal Pole Right |
| Frontal Pole Left | Frontal Pole Left |
| Lingual Gyrus Right | Lingual Gyrus Right |
| Lingual Gyrus Left | Lingual Gyrus Left |
| Occipital Pole Right | Occipital Pole Right |
| Occipital Pole Left | Occipital Pole Left |
| Precuneus Right | Precuneus Right |
| Precuneus Left | Precuneus Left |
| Parahippocampal Gyrus Right | Parahippocampal Gyrus Right |
| Parahippocampal Gyrus Left | Parahippocampal Gyrus Left |
| Planumtemporale Right | Planumtemporale Right |
| Planumtemporale Left | Planumtemporale Left |
| Supplementary Motor Cortex Right | Supplementary Motor Cortex Right |
| Supplementary Motor Cortex Left | Supplementary Motor Cortex Left |
| Supramarginal Gyrus Right | Supramarginal Gyrus Right |
| Supramarginal Gyrus Left | Supramarginal Gyrus Left |
| Superiorparietal Lobule Right | Superiorparietal Lobule Right |
| Superiorparietal Lobule Left | Superiorparietal Lobule Left |
| Temporal Pole Right | Temporal Pole Right |
| Temporal Pole Left | Temporal Pole Left |
| Temporal Gyrus Right | Right Inferior Temporal Gyrus; Right Medial Temporal Gyrus; Right Planum Polar; Right Superior Temporal Gyrus; Right Transverse Temporal Gyrus |
| Temporal Gyrus Left | Left Inferior Temporal Gyrus; Left Medial Temporal Gyrus; Left Planum Polar; Left Superior Temporal Gyrus; Left Transverse Temporal Gyrus |
| Orbital Gyrus Right | Right Anteriororbital Gyrus; Right Gyrus Rectus; Right Lateral Orbital Gyrus; Right Medial Frontal Cortex; Right Medial Orbital Gyrus; Right Posterior Orbital Gyrus; Right Subcolossal Area |
| Orbital Gyrus Left | Left Anteriororbital Gyrus; Left Gyrus Rectus; Left Lateral Orbital Gyrus; Left Medial Frontal Cortex; Left Medial Orbital Gyrus; Left Posterior Orbital Gyrus; Left Subcolossal Area |
| Cingulate Gyrus Right | Right Anterior Cingulate Gyrus; Right Middle Cingulate Gyrus; Right Posterior Cingulate Gyrus |
| Cingulate Gyrus Left | Left Anterior Cingulate Gyrus; Left Middle Cingulate Gyrus; Left Posterior Cingulate Gyrus |
| Frontal Gyrus Right | Right Superior Frontal Gyrus; Right Superior Frontal Gyrus Medial Segment; Middle Frontal Gyrus |
| Frontal Gyrus Left | Left Superior Frontal Gyrus; Left Superior Frontal Gyrus Medial Segment; Middle Frontal Gyrus |
| Occipital Gyrus Right | Right Superior Occipital Gyrus; Right Inferior Occipital Gyrus; Right Middle OccipitalGyrus |
| Occipital Gyrus Left | Left Superior Occipital Gyrus; Left Inferior Occipital Gyrus; Left Middle OccipitalGyrus |
| Inferior Frontal Gyrus Right | Right Tringular Part Of The Inferior Frontal Gyrus; Right Orbital Part Of The Inferior Frontal Gyrus; Right Opercular Part Of The Inferior Frontal Gyrus |
| Inferior Frontal Gyrus Left | Left Tringular Part Of The Inferior Frontal Gyrus; Left Orbital Part Of The Inferior Frontal Gyrus; Left Opercular Part Of The Inferior Frontal Gyrus |
| Operculum Right | Right Central Operculum; Right Frontal Operculum; Right Parietal Operculum |
| Operculum Left | Left Central Operculum; Left Frontal Operculum; Left Parietal Operculum |
| Insula Right | Right Posterior Insular; Right Anterior Insula |
| Insula Left | Left Posterior Insular; Left Anterior Insula |
| Postcentral Gyrus Right | Post Central Gyrus Right; Right Postcentral Gyrus Medial Segment |
| Postcentral Gyrus Left | Post Central Gyrus Left; Left Postcentral Gyrus Medial Segment |
| Precentral Gyrus Right | Precentral Gyrus Right; Right Precentral Gyrus Medial Segment |
| Precentral Gyrus Left | Precentral Gyrus Left; Left Precentral Gyrus Medial Segment |
| Fusiform Gyrus Right | Right Fusiform; Right OccipitalFusiform Gyrus |
| Fusiform Gyrus Left | Left Fusiform; Left OccipitalFusiform Gyrus |

**Supplementary Notes on Modelling**

***Dynamical systems model for HD disease progression***

We apply a previously established framework via adaptation to our HD progression sample. We summarize the relevant components of the model here and refer the mathematically interested reader to a more technical introduction of inference methods^1^. The dynamical system used for modelling brain changes is described via state model

$$\frac{dx}{dt}(t)=Ax(t)+Cu\left( t,\theta_{u} \right)$$

and observational (or measurement) model

$$y\left( t \right)=g\left( x(t),\theta_{g} \right)+\epsilon$$

with multivariate observations $y(t)$, state variables $x(t)$*,* system inputs $u(t,.)$, connectivity parameter matrix $A$, regional sensitivity parameter to inputs $C$, and residuals $\epsilon$*.*

Observations were available at discrete time-points with MRI acquisitions. In this application the model refers to states as being regional brain volumes within HD patients. More specifically, the state equation describes the temporal progression of 27 bilateral volumes (25 cortical regions, as well as the caudate and putamen) and one global WM volume over up to 10 years. In summary, the main generative model makes predictions for 55 brain regions using 28 dynamical state variables (illustrated in Supplementary Figure 2). The progression of states is influenced by both, endogenous dynamics $Ax(t)$, and external time-varying inputs $u(t,\theta_{u})$ with optional input parameters $\theta_{u}$. In the main findings (presented in Figures 1-3) the endogenous dynamics of the HD model were restricted to regional self-connections, i.e. *A* is a diagonal matrix. The diagonal elements can be interpreted as region-specific atrophy (or decay) rates $a_{ii}=a_{0}e^{\lambda_{i}}$ (with $a_{0}=-0.005;$i.e. using log-normal priors for enforcing negativity), causing decay which results (for our value range) in approximately linear volume loss over course of the progression.

It is important to mention that in our model we assumed bilateral symmetry of disease progression across hemispheres and thus the very same state variable describes the evolution of volumes in both corresponding bilateral grey matter ROIs (via using a linear observational model that averages both hemispheres observations to estimate one state). This symmetry is based on previous sMRI results reporting largely symmetric effects of atrophy in the TRACK-HD cohort and in a meta-analysis of HD studies^2–4^ and is a common practice in Structural Equation Modelling^5^ since rendering brain state variables estimates of latent (i.e. *error-free*) variables. Using this framework, the analyses presented above were addressed to measure gross atrophy, approximate linear rate of atrophy (via decay rate), potential non-linear accelerations of atrophy and the effects of CAG length regional decay rates.

***Exploring different inputs and alternative models***

External system inputs are additional drivers for volumetric state changes. In this model they describe unknown underlying factors that can influence atrophy within a region on top of its endogenous decay. This can be used to implement explicitly defined external input factors that affect regions differentially through estimated input sensitivity (or amplitude) parameters *C*. The following inputs (illustrated in Supplementary Figure 2B) were implemented:

(a) Linear velocity model (i.e. rate of atrophy is a linear function of time);

(b) Piecewise linear progression assuming a global acceleration phase;

(c) Velocities progressing in a sigmoidal (s-shaped) manner. These accelerations can have regionally-specific sensitivity to change and rates of change, but with a global delay parameter common across all regions reflecting a delay in the acceleration of atrophy. The global delay parameter could be before or after motor diagnosis;

(d) Velocities progressing in a sigmoidal (s shaped) manner. These accelerations can have regionally-specific sensitivity to change and rates of change, but this time with a regional delay parameter that reflects the possibility of a differing delay in the acceleration of atrophy across different regions. Again, the regional delay parameters could be before or after motor diagnosis;

(e) State equations follows a generalized logistic diff. equation (Richards curve);

(f) Volume change follows a simple quadratic polynomial progression

(g) Volumetric change evolves without the effects of external inputs

More specifically, to account for a potential acceleration of disease pathology we studied sigmoidal inputs

$$Cu\left( t,\theta_{u} \right)= -c/{(1+e^{-b(t-m)})}$$

with global or regional amplitudes $c$, time-shifts $m$, and (always global) rate of change parameter $b$ (sigmoid illustrated in Figure 2B). The sigmoidal amplitude parameter (shown in Figure 2B) allowed for regional acceleration of shrinkage during progression ($c>0$), decelerations ($c<0$), or no contribution of the input ($c=0)$. The time-shift parameter (shown in Figure 2C) of each regional sigmoid enabled earlier ($m<0)$or later accelerations ($m>0)$ in years relative to motor diagnosis.

Bayesian model comparisons revealed highest evidence for sigmoidal progression models with additional inputs (Supplementary Figure 2C) causing non-linearities with regional variations of both, the (1) temporal time-shift of accelerations/decelerations and (2) the sensitivity or amplitude of the accelerations We present the winning model group results in terms of the posterior distribution (expectation ± SD) of the group mean parameters in Fig. 1-3.

***Individual and*** ***group level inference***

AlI above first level model inversions were performed using previously established Variational Laplace methods^6^. More specifically, system integration and estimation followed a symmetric scheme from time point of motor diagnosis (treated as the initial volume state $x\left( t_{0} \right)$of the system since data was available for all participants) both forward and backwards in time to the earliest time point prior to diagnosis and the latest time point after motor diagnosis.

The aim of the study is a group level disease progression model, fully accounting and potentially explaining individual level non-linear trajectories. All patients first level models were embedded in a second (group-) level model. Here we took advantage of the a recently introduced Parametric Empirical Bayes (PEB) framework for estimation and inference on hierarchical non-linear models^7^. As in Ziegler et al.^1^, weakly informative priors were used for first level and second level parameters to allow results better reflect aspects of data rather than strong prior knowledge. Patient’s subject-specific characteristics such as cag-gene repeat length, sex, age at motor diagnosis, total intra-cranial volume and scanning site were included as covariates in the group level explaining first level variability.

Notably, since CAG repeat length and age at diagnosis were found to be correlated highly (r=-0.85), age was entered after orthogonalization with respect to cag. The hierachical modelling (a) accounts for undesired variation of first level parameters; (b) explicitly allows assessing effects of e.g. cag gene repeat length on all model parameters (e.g. decay rates); and (c) increases power for group level effects by accounting for first level parameter uncertainty differences across patients (e.g. scanned 3 vs. 7 times). Bayesian model selection (e.g. across inputs above) was conducted comparing the obtained full hierarchical (two-level) models with all described first level forms and a second level including a group mean parameter and above mentioned individual covariates and confounds. Bayesian model evidence accounts for both, optimizing model fit and while penalizing complexity and is therefore suitable for model selection in highly parameterized hierarchical disease progression models^8^.

***Predicting motor symptom changes using hidden brain states***

Main findings (presented in Figure 1-3) were restricted to analysis of disease progression dynamics of atrophy in 55 brain regions based on 28 state variables. In particular, that means that observational model predicts only brain observations. However, we finally extended the observational model to additionally predict motor deficit scores (via regression) using a linear combination of brain states:

$$y_{tms}(t)=w_{0}+\sum_{i=1}^{28} w_{i}x_{i}(t)+\varepsilon$$

with regional weights $w_{i}$describing the contribution of each regional atrophy state to the prediction of the individual motor score available for all brain scans in 47 subjects. Notably, the comparably large number of observations per subject in this exceptional dataset allows assessing the association of brain states and motor scores over time-points within-subject, in contrast to many conventional between-subject brain-behavioural findings.
